# NOTE ON A SMALL FRUIT-EATING BAT FROM THE MIDDLE HOLOCENE OF LAGOA SANTA, EASTERN BRAZIL

**DOI:** 10.1101/2024.12.13.628429

**Authors:** Artur Chahud

## Abstract

Bats represent the second most diverse order of mammals; however, their fossil and subfossil remains are still relatively understudied. In Brazil, numerous living species have been identified in cave deposits, although few fossils have been dated, with none originating from the Holocene. This study presents a specimen from the Middle Holocene, collected from Cuvieri Cave. The fossil consists of an incomplete, U-shaped mandible that includes the first molars. The specimen was identified as *Artibeus* sp., a genus of fruit-eating bats that is currently widespread across Brazilian territory. This specimen dates to a poorly understood period in the Lagoa Santa region, when human populations had abandoned the area (referred to as the Archaic Gap) and during which the faunal population of Cervidae experienced a notable decline in the number of fossil records. The genus *Artibeus* is widely distributed in tropical regions and is commonly found in environments ranging from semi-arid regions to savannas and dense forests. It is not considered a paleoenvironmental indicator.

## INTRODUCTION

The Holocene is the second epoch of the Quaternary period and encompasses the current era of Earth’s geological history. It began approximately 11,650 years ago, following the last glacial period and the conclusion of the Pleistocene. The Holocene is traditionally divided into three stages: the Greenlandian (11,650 to 8,326 years ago), the Northgrippian (8,326 to 4,200 years ago), and the Meghalayan (from 4,200 years ago to the present).

During the Late Pleistocene and Early Holocene in Brazil, the modern fauna became established, while mammalian megafauna, including giant sloths, saber-toothed cats, and glyptodonts, went extinct (PAULA COUTO, 1979; CHAHUD, 2020a; CHAHUD et al. 2023; 2024b).

In the Lagoa Santa region, the transition from the Early to the Middle Holocene was characterized by climatic changes and the disappearance of evidence of human communities in the area. This period, referred to as the Archaic Gap (ARAUJO et al. 2005; 2012; 2018), persisted for much of the Middle Holocene, remains poorly understood, and lacks comprehensive studies, particularly regarding its faunal record.

Chiroptera (bats) represent the second most diverse order of mammals, inhabiting a wide range of environments and exhibiting diverse ecological lifestyles. They are the only mammals capable of true flight. Despite their evolutionary and ecological significance, their paleontological history remains poorly understood, likely due to their small body size, behavioral traits, and fragile bone structure (PAULA COUTO, 1979). Geological records of bats include species from the Eocene in the Northern Hemisphere (TABUCE et al., 2019), though in South America, the most common bat specimens date from the Quaternary.

Specimens discovered in cave deposits are common and are typically associated with living species (CZAPLEWSKI & CARTELLE, 1998; FRACASSO & SALLES, 2005). These remains are usually represented by fragmented phalanges and metapodials, while fragments of mandibles or skulls are relatively rare. In the Lagoa Santa region, WINGE (1893) identified skeletal remains associated with extant Chiroptera species in local cave sites.

It is noteworthy that only two specimens discovered in Brazilian caves have been dated (CZAPLEWSKI & CARTELLE, 1998), both originating from the end of the Pleistocene. One of these was indirectly dated through analysis of an associated coprolite. However, many of these specimens result from bone remobilization, implying that they could be relatively recent, including both living and extinct species. *Desmodus draculae*, the only extinct Chiroptera species in Brazil and identified from late Pleistocene deposits, may have gone extinct more recently, during the late Holocene (PARDIÑAS & TONNI, 2000).

The Cuvieri Cave, located in the Lagoa Santa region, contains paleontological material that has undergone minimal remobilization. Some of these specimens have been dated and collected in accordance with the local stratigraphy, allowing for the establishment of age associations for various specimens. A small mandible belonging to a Stenodermatinae was collected from layers dated to the Middle Holocene. The objective of this study is to present this specimen, emphasizing its morphological and anatomical characteristics.

## MATERIAL AND METHODS

The Cuvieri Cave (Figure 1), located in the municipality of Matozinhos, in the Lagoa Santa region, Minas Gerais, is part of the Lagoa Santa Karst Environmental Protection Area and is located at UTM coordinates 23K 603756E and 7846105S (HUBBE et al., 2011). The cave entrance is approximately 1.5 meters high by 1 meter wide, giving access to a main conduit that is 15 meters long. This passage has three independent vertical shafts, called *Locus* 1, *Locus* 2 and *Locus* 3 (Figure 1), with depths of 16, 4 and 8 meters, respectively (HUBBE et al., 2011). The specimen was collected from *Locus* 2.

**Figure 1.**
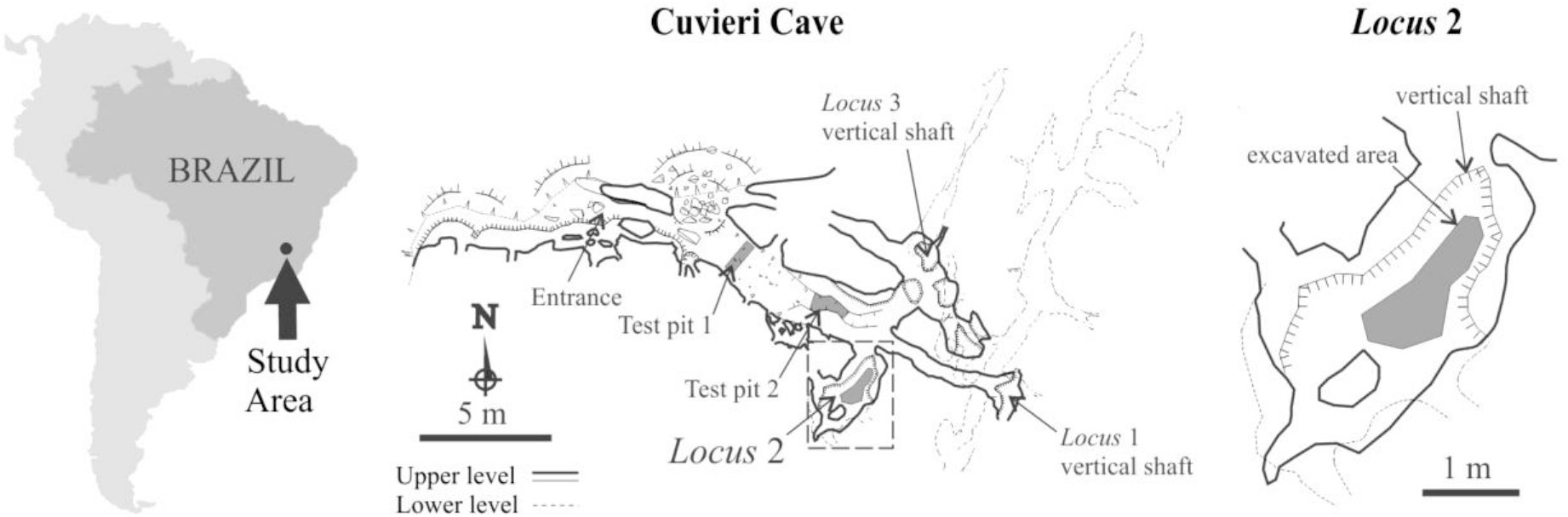
Geographic location of the study area and of Cuvieri Cave showing the position of *Loci* 1, 2 and 3 (map courtesy of Alex Hubbe and Grupo Bambuí for Speleological Research).

The cave features an accumulation of poorly consolidated terrigenous sediment, which allowed for excavations with rigorous stratigraphic control. According to HUBBE et al. (2011), the excavation techniques involved exposing depositional surfaces, resulting in the identification of 46 exposures in *Locus* 2.

The identification of the specimen was made based on comparison with known specimens and consulted the works of WEBSTER & JONES (1982a; 1982b), TIMM (1985), FREEMAN (1988), ORTEGA & CASTRO-ARELLANO (2001), LIM et al. (2004), HOLLIS (2005), ZORTÉA & TOMAZ (2006), CARVALHO et al. (2014), LÓPEZ-AGUIRRE & PÉREZ-TORRES (2015), MORALES-MARTÍNEZ & RAMÍREZ-CHAVES (2015), ORTEGA et al. (2015), VILAR et al. (2016), GARBINO & TAVARES (2018), SALAS et al. (2018), PALACIOS-MOSQUERA et al. (2019), BRANDÃO & HINGST-ZAHER (2021), CZAPLEWSKI & BAKER (2022), VRLA et al. (2023).

The specimen analyzed in this study comes from Exposure 21 of *Locus* 2, which also contained a specimen of *Subulo cf. gouazoubira* dated to 7050 ± 50 years BP (Before Present), a period falling within the Middle Holocene (Northgrippian).

As a small and highly fragile bone fragment, it is likely that the specimen was rapidly buried with minimal remobilization or reworking. The primary alteration it underwent was the loss of most of its dentition due to disarticulation, with the alveoli remaining almost intact. This suggests that the specimen was unlikely to have been subject to significant hydraulic reworking from older layers, implying that the fragment likely originates from the Middle Holocene.

The specimen was cataloged under the registry number CVL2EXP21-1 and is curated in the Laboratory of Human Evolutionary Studies of the Institute of Biosciences of the University of São Paulo (LEEH-IB-USP). The bone piece was collected by sieving during sampling and received the acronym “CVL2” (Cuvieri *Locus* 2), indicating the place of its discovery, followed by “EXP21” (specific exposure) and the corresponding registry number.

## SYSTEMATIC PALEONTOLOGY

Order: Chiroptera Blumenbach, 1779

Family: Phyllostomidae Gray, 1825

Subfamily Stenodermatinae Gervais, 1856

*Artibeus* sp.

Figure 2

**Figure 2.**
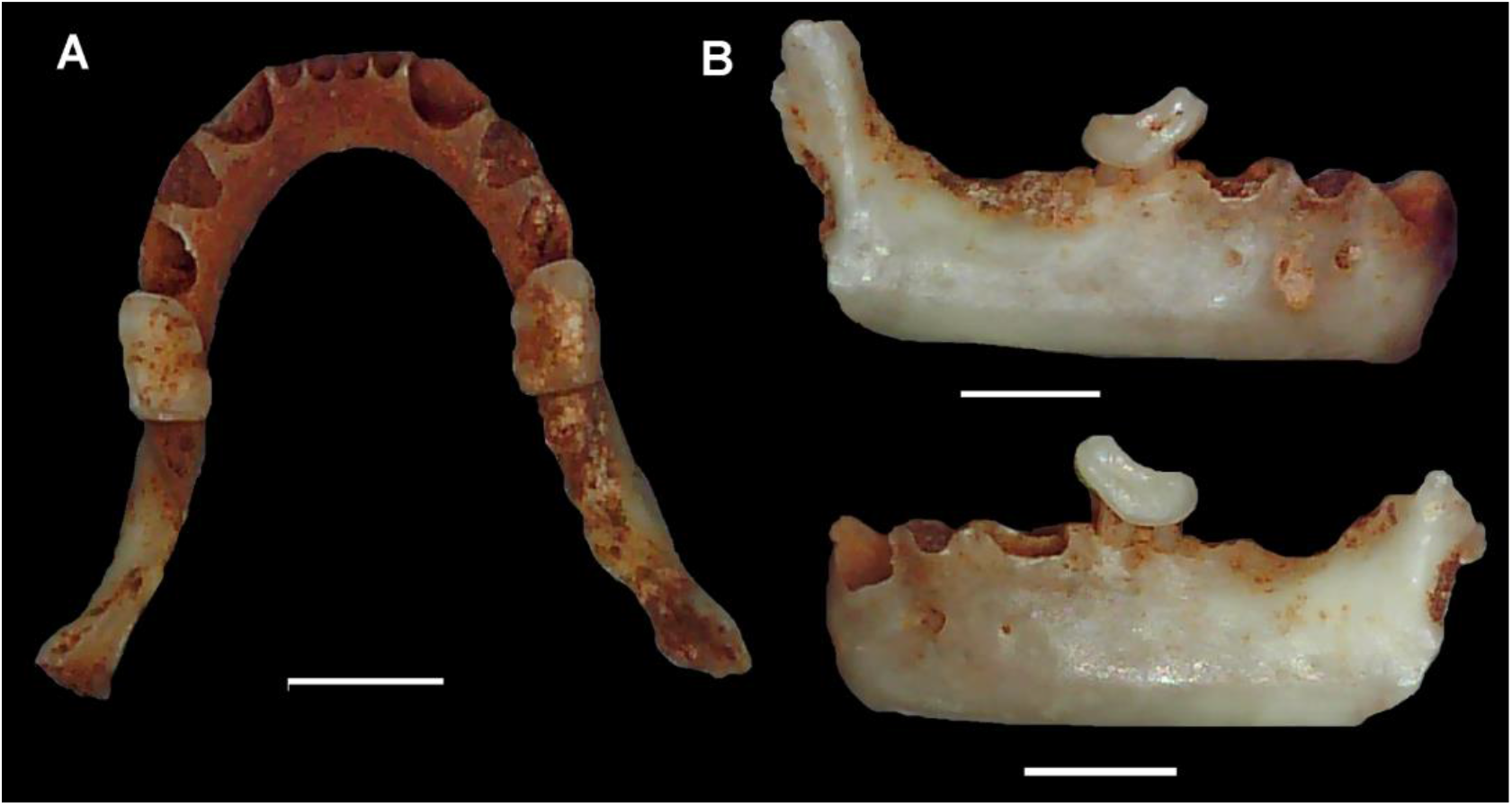
Mandible of the specimen found in Cuvieri Cave, CVL2EXP21-1. A) dorsal view; B) lateral views. Scale 2mm.

### Material

Incomplete mandible, containing only the first two molars (M1) of opposite positions (CVL2EXP21-1).

### Remarks

The specimen consists of a partially fragmented mandible, with dimensions of 8.9 mm in length and 4.7 mm in height. The alveoli are well-preserved, allowing for the observation that the diameter of the incisors is considerably smaller than that of the canines, premolars, and molars (Figure 2A).

The angle formed by the front curvature of the mandible, separating the two sides, is approximately 60°. The intersection between the sides of the mandible presents a “U” shape, with the side almost parallel to each other, except for a slight divergence in the articulatory region (Figure 2A).

The preserved molar teeth are 1.5 mm in length in the anteroposterior direction and approximately 1 mm in width in the lingual-labial direction.

The anterior portion of the molar teeth is more prominent than the posterior portion (Figure 2B), a common characteristic in fruit-eating bats. Furthermore, the molars do not exhibit prominent cusps, unlike other species of Phyllostomidae (GARBINO & TAVARES, 2018). The dental crown does not show carnassiform characteristics, nor are there any prominent ridges typically found in insectivorous bats (CZAPLEWSKI & BAKER, 2022).

## Discussion

The observed molar differs from those found in omnivorous, insectivorous, nectarivorous, and carnivorous bats due to the absence of carnassiform features and the general “U”-shaped structure of the mandible (MORALES-MARTÍNEZ & RAMÍREZ-CHAVES, 2015; BRANDÃO & HINGST-ZAHER, 2021; CZAPLEWSKI & BAKER, 2022). Furthermore, the position and characteristics of the preserved alveoli, as well as the presence of the large molar, contrast with the features observed in hematophagous bats, such as *Desmodus*.

The angle formed by the frontal curvature of the mandible supports the hypothesis that the specimen belongs to a fruit-eating bat, since animalivorous bats have more elongated maxillae and mandibles, with angles below 50° (FREEMAN, 1988).

When compared to other genera of fruit-eating bats, such as *Vampyrodes*, the specimen from the Cuvieri Cave displays a “U”-shaped mandible (CARVALHO et al., 2014). However, the incisors and molar of this specimen are larger and more prominent than those in *Vampyrodes*.

The morphology of the mandible and the preserved dentition are conpatible with those found in species of the genera *Artibeus* and *Dermanura* (WEBSTER & JONES, 1982a; 1982b; TIMM, 1985; ORTEGA & CASTRO-ARELLANO, 2001; HOLLIS, 2005; ORTEGA et al., 2015), both of which belong to the subfamily Stenodermatinae. According to LIM et al. (2004), no diagnostic features clearly distinguish *Dermanura* from *Artibeus*. However, size differences and discrete molecular lineages are recognizable, leading the authors to suggest that *Dermanura* should be considered a subgenus within *Artibeus*. In the present study, we have accepted *Artibeus* as the valid genus, with *Dermanura* regarded as a subgenus, as proposed by LIM et al. (2004).

When comparing with other specimens of *Artibeus*, the more slender mandible morphology and the significant height relative to the total preserved size do not allow an association with any known species within this genus.

The dentition of various species within the genus *Artibeus* and the subgenus *Dermanura* features a first molar similar to that of the specimen found, as well as prominent incisors (PALACIOS-MOSQUERA et al., 2019). However, this molar may also exhibit prominent cusps in some species, as observed in *A. fraterculus* (SALAS et al., 2018).

It is important to note that cranial and mandibular abnormalities and asymmetries have been observed in individuals of *Artibeus lituratus*, suggesting the occurrence of individual variations within the species (LÓPEZ-AGUIRRE & PÉREZ-TORRES, 2015; VRLA et al., 2023). This characteristic makes identification at the species level difficult and, therefore, only the generic identification was chosen for the specimen, *Artibeus* sp.

Bats of the genus *Artibeus* are found in several Brazilian biomes, inhabiting environments such as the semi-arid region, savanna and dense forests (ZORTÉA & TOMAZ, 2006; VILAR et al., 2016; BRANDÃO & HINGST-ZAHER, 2021). However, this genus has not been recorded in temperate or cold climates, being characteristic of tropical regions.

## ASSOCIATED FAUNA

The period between 8,500 and 6,000 years BP, corresponding to the Middle Holocene, is poorly understood with regard to the paleofauna of the Cuvieri Cave. Observations by CHAHUD & OKUMURA (2023) and CHAHUD et al. (2024a) identified the presence of the cervid *Subulo gouazoubira* during this period.

During the Pleistocene, the Early Holocene, and the periods more recent than 6,000 years BP, the fauna of the Cuvieri Cave is characterized by the presence of Leporidae, Anura, Tapiridae, the cervid *Mazama americana*, and the tayassuid *Dicotyles tajacu* (CHAHUD et al., 2020; CHAHUD, 2020b, 2021, 2022; CHAHUD & OKUMURA, 2021, 2022, 2023). However, the records of these vertebrate groups have not been robustly confirmed or are represented by few bones, generally fragmented, found in strata dated to the Middle Holocene, suggesting the remobilization of deeper layers.

The material that is either minimally or not remobilized, free from breakage or signs of abrasion, in addition to *Subulo gouazoubira*, includes a few appendicular bones of small rodents, marsupial teeth, and Squamata bones. A study of *S. gouazoubira* in the Middle Holocene indicated a decline in the number of bones of this cervid at Locus 2 of the Cuvieri Cave (CHAHUD et al. 2024a), suggesting a slight population decrease during this period.

This period of faunal population decline coincided with what archaeologists have termed the Archaic Gap, due to the sparse or nonexistent evidence of human activity in archaeological sites in the Lagoa Santa region.

Paleoenvironmental studies suggest that the beginning of the Holocene was marked by a humid climate, followed by a dry period that likely began around 8,000 years BP and ended about 6,200 years BP (ARAUJO et al., 2005; 2012; 2018; BARROS et al., 2011). The hypothesis of increased humidity in the region between 6,000 and 5,000 years BP was proposed by PARIZZI et al. (1998), who indicated a significant increase in humidity from 5,400 years BP. This increase in humidity between 6,000 and 5,000 years BP is consistent with the increased amount of osteological material found in the Cuvieri Cave (CHAHUD et al., 2020; CHAHUD, 2022; CHAHUD & OKUMURA, 2023).

The Archaic Gap period has been a subject of debate, with RACZKA et al. (2013) challenging the hypothesis of aridity through palynological studies, which suggest that climatic instabilities affected the region. However, the absence or reduction in the record of several taxa and the slight population decline of *Subulo gouazoubira* observed in the Middle Holocene deposits of Cuvieri Cave (CHAHUD et al., 2024a) suggest that the environmental changes were not extreme. It is likely that the region experienced slightly drier conditions, but not drastically different from many regions of the current biome.

The presence of fruit-eating bats in the Lagoa Santa region during the Middle Holocene indicates that the climate was not excessively dry, as the vegetation necessary to support a biota specialized in fruit consumption was still present.

## CONCLUSIONS

The mandible discovered belongs to a small bat of the genus *Artibeus*. However, due to the partial preservation, with only the first two molars present, and the individual variations observed within species of this genus, it was not possible to securely determine the species.

Bats of the genus *Artibeus* are generalists in their consumption of plant material and, therefore, do not serve as good paleoenvironmental indicators. However, their presence suggests that there was sufficient vegetation in the Lagoa Santa region during the Middle Holocene, indicating that an excessively arid environment or extremes in humidity and temperature were likely uncommon during this period. This hypothesis suggests that the specimen probably inhabited an environment with vegetation similar to forests or shrublands, not significantly different from the current environment.

## ACKNOWLEDGEMENTS

The author thanks to Steve Langley for his English review, and to Dr. Maria Mercedes Martinez Okumura, responsible for LEEH (Laboratory for Human Evolutionary Studies), Department of Genetics and Evolutionary Biology, Institute of Biosciences of the University of São Paulo for permitting the preparation of the fossils in her laboratory.

